# Molecular analysis of long non-coding RNA GAS5 and microRNA-34a expression signature in common solid tumors: A pilot study

**DOI:** 10.1101/325175

**Authors:** Eman A. Toraih, Saleh Ali Alghamdi, Aya El-Wazir, Marwa M Hosny, Mohammad H. Hussein, Motaz S. Khashana, Manal S. Fawzy

## Abstract

Accumulating evidence indicates that non-coding RNAs including microRNAs (miRs) and long non-coding RNAs (lncRNAs) are aberrantly expressed in cancer, providing promising biomarkers for diagnosis, prognosis and/or therapeutic targets. We aimed in the current work to quantify the expression profile of miR-34a and one of its bioinformatically selected partner lncRNA growth arrest-specific 5 (GAS5) in a sample of Egyptian cancer patients, including three prevalent types of cancer in our region; renal cell carcinoma (RCC), hepatocellular carcinoma (HCC) and glioblastoma (GB) as well as to correlate these expression profiles with the available clinicopathological data in an attempt to clarify their roles in cancer. Quantitative real-time polymerase chain reaction analysis was applied. Different bioinformatics databases were searched to confirm the potential miRNAs-lncRNA interactions of the selected ncRNAs in cancer pathogenesis. GAS5 was significantly under-expressed in the three types of cancer. However, levels of miR-34a greatly varied according to the tumor type; it displayed an increased expression in RCC [4.05 (1.003-22.69), *p* <0.001] and a decreased expression in GB [0.35 (0.04-0.95), *p* <0.001]. A weak negative correlation was observed between levels of GAS5 and miR-34a in GB [r = −0.39, *p* =0.006]. Univariate analyses revealed a correlation of *GAS5* downregulation with poor disease-free survival (r = 0.31, *p* =0.018) and overall survival (r = 0.28, *p* =0.029) in RCC but not in GB, and a marginal significance correlation with a higher number of lesions in HCC. Hierarchical clustering analysis showed RCC patients among others, could be clustered by GAS5 and miR-34a co-expression profile. Our results confirm the tumor suppressor role of GAS5 in cancer and suggest its potential applicability to be a predictor of bad outcomes with other conventional markers for various types of cancer. Further functional validation studies are warranted to confirm miR-34a/GAS5 interplay in cancer.

## Introduction

Cancer is now the second leading cause of mortality worldwide, causing 8.8 million deaths globally in 2015, which is equivalent to one in every six deaths [1]. Cancer occurs as a net result of activation of oncogenes and inhibition of tumor suppressor genes (TSGs) [2]. Decades back, it was believed that oncogenes and TSGs had to code for proteins, and only mutations in protein-coding genes would result in such pathological conditions as cancer. However, in the world of genetics, it is never that simple. With the advancement of genetic technologies including next generation sequencing, microarrays and bioinformatic machinery, many truths came to light, including what was originally thought to be junk DNA is now found to code for thousands of equally significant regulatory RNAs [3]. Since their discovery, the non-coding RNAs (ncRNAs) have been recognized as epigenetic regulators of protein-coding genes. Recently, a whole new level of regulation has been uncovered, when it was found that ncRNAs have the ability to regulate each other as well [4] further adding to the complexity of the regulatory processes. This set of findings has revolutionized our understanding of several human diseases, making this ‘the era of non-coding RNAs’.

Many classes of ncRNAs have been identified and linked to cancer, the most common of which are micro RNAs (miRNAs), long-noncoding RNAs (lncRNAs), PIWI-interacting RNAs (piRNAs) and small nucleolar RNAs (snoRNAs). The roles of these four types of ncRNAs in cancer are reviewed in several studies [5–8]. In brief, overexpression of some ncRNAs as miRNAs or lncRNAs can suppress the expression of TSG targets, while loss of function or reduced expression of others may allow overexpression of the oncogenes they regulate. Furthermore, since each ncRNA may regulate hundreds of different genes, its over or under expression may have widespread oncogenic effects because many genes will be dysregulated [9].

Here we were interested in a relatively newly discovered lncRNA; Growth Arrest Specific 5 (GAS5) which is a poorly conserved gene mapped to chromosome 1q25.1 [10]. It consists of 12 exons and 11 introns from which 29 transcripts are produced from alternative splicing, many of which contain retained introns (Ensembl Genome Browser ’GAS5’). The first two transcripts discovered (produced from alternative splice sites on exon 7) are the GAS5a and GAS5b with the latter being predominantly expressed in most cells [11]. Another synonym for the gene is’ Small Nucleolar RNA Host Gene’, this is for having the ability to produce multiple (10 in human) noncoding small nucleolar RNAs (snRNAs) from its more conserved introns [11]. These snRNAs are involved in the regulation of ribosomal RNA (rRNA) synthesis through 2-O-methylation of pre-ribosomal RNA [12].

As its name implies, GAS5 is over expressed in growth-arrested cells [13]. This is further demonstrated by the presence of high levels of GAS5 in brain cells, which are considered the slowest dividing cells in the body as opposed to its lowest levels in other rapidly dividing cells, the most important of which are cancer cells [14].

Expectedly, GAS5 under expression was found to be associated with multiple types of cancer [15–25] and it was found to bind some miRNAs, including miR-21, miR-222 and miR-103 sponging their inhibitory effect on their target TSGs ([16, 21, 26, 27].

Our bioinformatic analyses have revealed a potential new miRNA target for the GAS5 gene; miR-34a. This miRNA is encoded from chromosome1p36.22. Its promotor is recognized for having multiple CpG islands and a p53 binding site, making p53 a direct transcriptional regulator for this miRNA [28, 29]. While the history of miR-34a with cancer is very well established, studies conducted on different types of cancers have contradictory results regarding its actual role in tumor progression. In other words, is it an oncogene or a TSG? Many studies have proved it’s functioning as a TSG in various types of cancer, including neuroblastoma [30], leukemia [31], pancreatic [32] and hepatocellular [33, 34] carcinoma, glioblastoma (GB) [35, 36], breast [37] lung [38] and colon [39] cancer. On the contrary, other studies have found that it functions as an oncogene through promoting tumorigenesis as in RCC [40, 41], papillary thyroid carcinoma (PTC) [42], colon [39] and uterine cancer [43]. It has been suggested that this discrepancy may be attributed to the tissue type and the miR-34a/p53 pathway involved [41].

To the best of our knowledge, no clinical studies were conducted to explore both GAS5 and miR-34a profiles in cancer patients. Hence, we were interested to investigate the expression profiles of these ncRNAs in three prevalent types of cancer in our region; renal cell carcinoma (RCC), hepatocellular carcinoma (HCC) and glioblastoma (GB) as well as to correlate these expression profiles with the available clinicopathological data in an attempt to clarify their roles in cancer.

## Materials and Methods

### Genomic characterization of GAS5 and miR-34a

Chromosomal localization, genomic sequence and structure analysis, subcellular localization, variant analysis, and folding pattern were retrieved from different online tools; including Ensembl (http://www.ensembl.org/), GeneCards for human gene database (http://www.genecards.org/), National Center for Biotechnology Information (NCBI) (ncbi.nlm.nih.gov/), COMPARTMENTS subcellular localization (https://compartments.jensenlab.org/Search), Database of Transcription Start Sites (DBTSS) version 10.0 (http://dbtss.hgc.jp/), KineFold (http://kinefold.curie.fr/), and MFold webserver (http://unafold.rna.albany.edu/?q=mfold/RNA-Folding-Form).

### Exploring miR-34 and GAS5 interactions

Identifying complementary regions between microRNA-34a and lncRNA GAS5 were demonstrated by several tools; RNA22 microRNA target detection (https://cm.jefferson.edu/rna22/) and DIANA-LncBase v2 (http://diana.imis.athena-innovation.gr/DianaTools/index.php).

### Functional enrichment analysis

Pathway enrichment analysis and gene ontology of microRNA-34a was performed by Diana-miRPath v3.0 (http://diana.imis.athena-innovation.gr/DianaTools/index.php) using its experimentally validated gene targets. Functional enrichment analysis of GAS5 was obtained from Ensembl and GeneCards databases to identify its biological function in cancer.

### Study population and sampling collection

In total, 230 samples were analyzed, including 60 formalin-fixed, paraffin-embedded (FFPE) RCC samples and their paired adjacent non-cancer tissues, 50 FFPE GB specimens and 10 non-cancer brain tissues as well as 30 HCC blood samples and 20 controls.

#### (a) RCC cohort

The archived FFPE renal samples were taken from sixty patients who underwent radical nephrectomy for a primary RCC and dating back for three years. All retrieved cases were archived in the Pathology laboratory of Mansoura Oncology Center, Mansoura, Egypt. No history of receiving neoadjuvant chemotherapy or radiotherapy prior to sampling. Clinicopathological data, including the survival were collected from patient medical records. Paired sixty cancer-free adjacent tissues were examined and sectioned to serve as controls for molecular analysis.

#### (b) GB cohort

Fifty glioblastoma patients and 10 non-cancer brain tissues collected from the archive of the Pathology Department, Mansoura University Hospitals, Egypt, dating back for three years were included in the current work. Detailed patients’ data were retrieved from their follow up records. They had GB grade 4, undergone surgical removal and had not received any treatment before sampling. They followed-up for more than 3 years.

#### (c) HCC cohort

Following our local hospital and medical ethical committee rules in liver tissue sampling prohibition from HCC patients, only blood samples were available. Thirty HCV-induced HCC and 20 matched controls from healthy blood bank donors were recruited in the study. Patients were obtained from the outpatient clinic of Tropical Medicine and Gastroenterology Department, Faculty of Medicine, Assuit, Egypt. They had typical imaging findings of liver cancer and elevated alpha fetoprotein (AFP). Patients underwent clinical and radiological assessment, confirmation of HCV by PCR, Barcelona-Clinic Liver Cancer (BCLC) staging, and Child-Turcotte-Pugh (CTP) scoring (44). Survival data for HCC patients were not available in patients’ medical records. Hence, these data were not included in the statistical analysis for HCC patients.

### Ethical approval

The study was conducted according to the ethical guidelines of the Declaration of Helsinki and approved by the Medical Research Ethics Committee of Suez Canal University.

### RNA extraction

Total RNA; including the small RNA, was isolated from either FFPE tissue sections (5 to 8-μm-thick) using the Qiagen miRNeasy FFPE Kit (*Qiagen, cat no 217504*) or serum using Qiagen miRNeasy serum/plasma Kit (*Qiagen, cat no 217184*) following the protocols supplied by the manufacturer. Concentration of RNA was determined using the NanoDrop ND-1000 spectrophotometer (*NanoDrop Tech., Inc. Wilmington, DE, USA*). Samples with a 260/280 nm absorbance ratio less than 1.8 were excluded.

### Reverse transcription reaction

Subsequently, RNA for lncRNA GAS5 was converted to complementary DNA (cDNA) in a T-Professional Basic, Biometra PCR System (*Biometra, Goettingen, Germany*) using high Capacity cDNA Reverse Transcription Kit (*Applied Biosystems, P/N 4368814*) with RT random primers as previously described [45].

Reverse transcription of miR-34a was specifically converted to cDNA using TaqMan™ MicroRNA Reverse Transcription kit (*P/N 4366596; Applied Biosystems, Foster City, CA, USA*) with 5x miRNA specific stem-loop primers as previously described in our prior publication [46]. Appropriate controls were included in each experiment.

### LncRNA GAS5 and microRNA-34a expression analyses

The Real-Time PCR reactions were performed in accordance with the Minimum Information for Publication of Quantitative Real-Time PCR Experiments (MIQE) guidelines [47]. The expression level of *GAS5* was assessed via SYBR Green qPCR analysis and normalized with GAPDH. The following primers were designed using Primer 3 software and checked by *in silico* PCR amplification of the University of California, Santa Cruz (UCSC) genome browser; for *GAS5*: Forward: 5′- CTTGCCTGGACCAGCTTAAT-3′; Reverse: 5′-CAAGCCGACTCTCCATACCT-3′, and for *GAPDH*: Forward: 5’- CGGATTTGGTCGTATTGGG-3’; Reverse: 5’-CTGGAAGATGGTGATGGGATT-3’. In brief, 10 μl of qPCR Green Master (*Jena Bioscience, Cat no. PCR-313L*), 0.6 μl (10 μM) forward and reverse primers, 8 μl PCR grade water, and 1 μl cDNA template were included in the reaction for *GAS5* and *GAPDH* SYBR Green assay analyses [46]. For microRNA quantification, a final volume of 20 μl was adjusted in duplicate, including 1.33 μl specific RT products, 2× TaqMan^®^ Universal PCR Master Mix with UNG (Applied Biosystems, P/N 4440043), and 20× of 1 μl specific TaqMan^®^ RNA assay for hsa-miR-34a-5p (Applied Biosystems, assay ID 000426) and Taqman^®^ Universal PCR master mix II, No UNG (2×) [48]. Two endogenous control assays were used; TATA box binding protein (TBP; assay ID Hs00427620_m1) in brain cancer and RNU6B (assay ID Hs001093) in liver and renal cancer [49, 50]. Appropriate negative and positive controls were used. The PCR for 96-well plates was carried out using StepOne™ Real-Time PCR System (Applied Biosystem) and incubated as follows: 95°C for 10 min followed by 45 cycles of 92°C for 15 seconds and 60°C for 1 minute. Ten percent randomly selected study samples were re-evaluated in separate runs for the study gene expressions to test the reproducibility of the qPCR which showed very close quantitative cycles value results and low standard deviations.

### Statistical analysis

R package (version 3.3.2) and Statistical Package for the Social Sciences (SPSS) for Windows software (version 22.0) were used for data analyses. Categorical variables were compared using the Chi-square (χ^2^) or Fisher’s exact tests where appropriate. Wilcoxon matched-pair signed-rank and Mann-Whitney U tests were used for tissues of renal/brain cancer and serum of liver cancer patients, respectively, to compare continuous variables. The correlation between miR-34a level and *GAS5* expressions was calculated by Spearman’s rank correlation analysis. A two-tailed *p*-value of < 0.05 was considered statistically significant. The receiver operating characteristic (ROC) curves were performed to get the best cutoff values of miRNA-34a for discriminating long and short survivors in cancer patients. The fold change of ncRNAs expressions in each patient relative to the control was calculated via Livak method based on the quantitative cycle (Cq) values with the following equation: relative quantity = 2^−ΔΔ*Cq*^; where ΔΔ*C*_*q*_ = (*C*_*q*_ ncRNA − *C*_*q*_ internal control)_cancer_ - (*C*_*q*_ ncRNA - *C*_*q*_ internal control)_control_ [51]. Univariate analysis for association between ncRNA expression profile and clinico-pathological features in cancer patients was run. The software package named PC-ORD ver. 6 [52] was employed to run different multivariate analyses for clustering analysis of patients according to clinico-pathological and molecular data.

## Results

### Genomic location of GAS5

Lnc-RNA GAS5 is also known as small nucleolar RNA host gene 2 (SNHG2) and non-protein coding RNA 30 (NCRNA00030). It is encoded by GAS5 gene on chromosome 1q25.1 (Genomic coordination at 1:173863901-173867987 at the negative strand according to human genome assembly GRCh38) (Fig. 1A). It consists of 12 exons, spanning 4.087 kb and encoding for 29 different alternative splice transcripts ranging in length from 242 to 1698 bp (Electronic Supplementary Table S1).

**Fig. 1.**
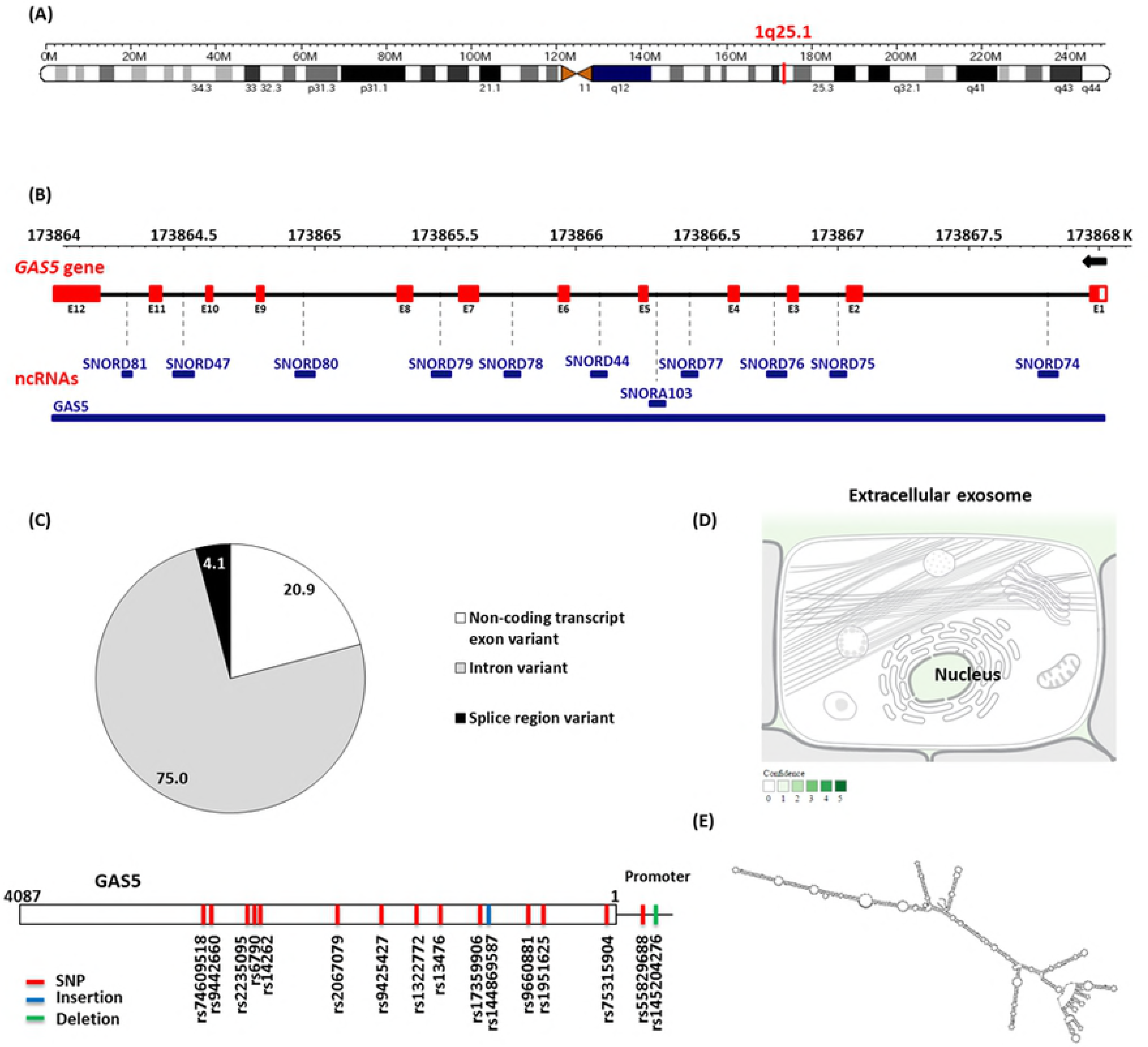
Structural analysis of *GAS5* gene. (A) Chromosomal localization of *GAS5* gene. It is present on chromosome 1q25.1, at genomic coordination 1:173863901-173867987 on the negative strand (according to human genome assembly GRCh38). (B) Sequence analysis of *GAS5* gene. It consists of 12 exons, spanning 4.087 kb that code for 29 different alternative splice transcripts. It hosts multiple snoRNA genes within its introns (except intron 9). These genes encode for ten C/D box snoRNAs, which contain the C (UGAUGA) and D (CUGA) box motifs, and an H/ACA box snoRNA, SNORDA103, within intron 4. (C) Genetic variant analysis. *GAS5* gene (4087 bases long) contains around 300 thousand variants (75% intronic, 21% exonic, and 4% splice region polymorphisms). Among all these polymorphisms, 14 SNPs (red), one deletion (green) and one insertion (blue) are common variants with minor allele frequency (MAF) > 0.05. (D) Subcellular localization of lncRNA GAS5. Text mining highlighted its predominant existence intranuclear and within extracellular exosomes which are extruded into the circulation. (F) Folding pattern of lncRNA GAS5. [Data source: Ensembl.org, genecards.org, NCBI, COMPARTMENT database, and MFold]

### Sequence analysis of *GAS5*

*GAS5* gene contains a seven-nucleotide oligo-pyrimidine tract on its 5′-end in exon 1, hence is classified as a member of the 5’ terminal oligo-pyrimidine (TOP) genes. This sequence can act as a cis-regulatory motif which either inhibits the binding of translational regulatory proteins downstream to the transcriptional start sites of mRNAs or suppresses the translational machinery itself. In addition to its translational controls, the TOP elements are known to modulate gene expression through regulating transcription [53]. Being one of the TOP genes, it is ubiquitously expressed and is predicted to regulate the translation of more than 20% of total mRNAs [DataBase of Transcription Start Sites]

Sequence analysis of *GAS5* gene revealed that it is a small nucleolar RNA host gene, containing multiple *snoRNA* genes within its introns. These genes encode for ten C/D box snoRNAs, which contain the C (UGAUGA) and D (CUGA) box motifs. They are predicted to play a role in the 2’-O-methylation of rRNA by guiding guanine methylation, which enhances RNA folding and interaction with ribosomal proteins. *GAS5* also hosts SNORDA103 within intron 4, an H/ACA box snoRNA, which is associated with pseudouridylation (Fig. 1B).

### Variant analysis

*GAS5* gene is shown to be highly polymorphic, enclosing around 300 thousand variants (75% intronic, 21% exonic, and 4% splice region polymorphisms). Among all these polymorphisms, 14 SNPs, one deletion and one insertion are common variants with minor allele frequency (MAF) > 0.05 (Fig. 1C) and (Electronic Supplementary Table S2).

### GAS5-miRNA interaction

Analysis with the RNA22 program (http://cbcsrv.watson.ibm.com/rna22.html) identified complementary regions of GAS5 with 690 microRNAs. Among these putative microRNAs, only 252 interactions with 234 microRNAs showed significant binding (p<0.05) (Electronic Supplementary Table S3). Via DIANA-LncBase v2 database for experimentally validated miRNA-lncRNA interactions, GAS5 was identified as a miR-34a target by immunoprecipitation experiments (score=0.558). For further confirmation, we used RNA22 software to determine the interaction binding sites between GAS5 transcripts and miR-34a. Our results showed base-pairing in twenty-three alternative splice variants. Among them, nine transcripts had two miR-34a binding sites (Electronic Supplementary Table S4).

### miR-34a functional analysis

Hundreds of miR-34a-5p and 3p targets were retrieved from various online databases; including miRTarBase (http://mirtarbase.mbc.nctu.edu.tw/), miRDB (http://mirdb.org/) and microRNA.org. Functional enrichment analysis revealed its enrollment in cancer-related KEGG pathways, as Pathways in cancer (hsa05200, 115 targets, p=0.001723304), proteoglycans in cancer (hsa05205, 71 targets, p=1.687731e-06), adherens junction (hsa04520, 34 targets, *p* = 3.396929e-06), cell cycle (hsa04110, 54 targets, *p* = 3.355808e-05), and p53 signaling pathway (hsa04115, 31 targets, *p* = 0.006060223) as well as cancer-specific pathways, namely glioma (hsa05214, 29 targets, *p* = 0.0003111577) and renal cell carcinoma (hsa05211, 28 targets, p= 0.02521455) (Electronic Supplementary Table S5). GO analysis (Diana tools) demonstrated miR-34a to be involved in cell death, cell cycle, and response to stress thus highlighting its role in cancer cell growth. In addition, miR-34a was significantly associated with membrane organization, cell junction organization, and cell motility, hence may play a key role in cancer cell invasion and metastasis (Fig. 2).

**Fig. 2.**
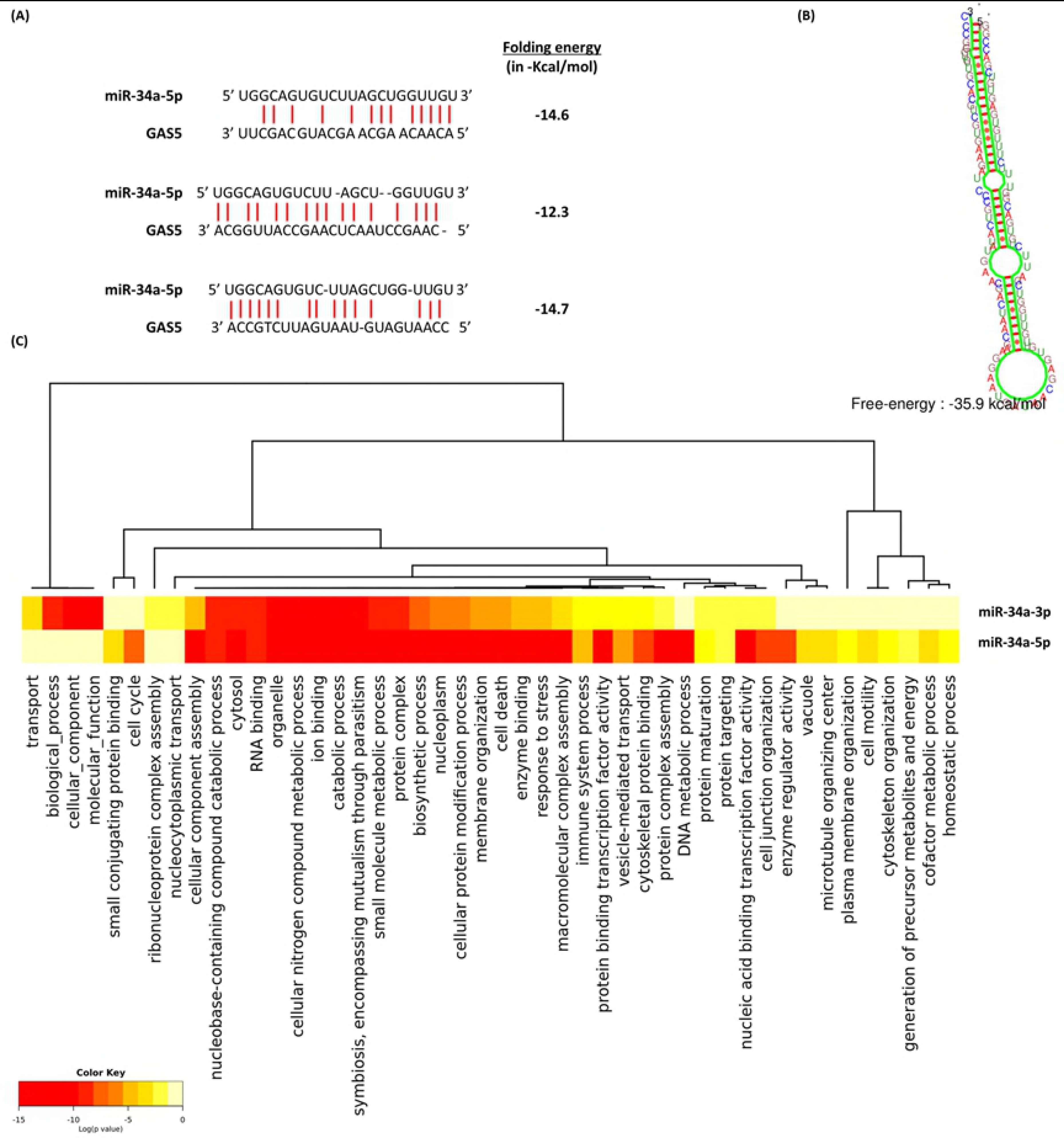
Functional and structural analysis of miR-34a. (A) GAS5: miR-34a-5p interaction. Complementarity regions are shown in three binding sites in *GAS5* gene, one proximal at 5’ region (1432-1455) and two distal at 3’ end (3545-3567 and 3698-3719) [Data source: RNA22, DIANA-LncBase v2]. (B) Predicted secondary structure of miR-34a. Folding pattern and energy are demonstrated [Data source: KineFold]. (C) Functional enrichment analysis of miR-34a experimentally validated gene targets. Significant clustered heat map represents the gene ontology using GO slim option, FDR conservative states, *p* value threshold < 0.05, and categories union [Data source: DIANA-miRPath v3]. http://snf-515788.vm.okeanos.grnet.gr/uploads/R/HeatMap290913.png

### Characteristics of the study population

Baseline characteristics of RCC, GB, and HCC cohorts are demonstrated in Tables 1 to 3.

**Table 1.** Clinicopathological characteristics of renal cell carcinoma patients.

| Variables | Total | Low OS | High OS | *p* value |
| --- | --- | --- | --- | --- |
| **Total number** | 60 (100) | 19 (31.7) | 41 (68.3) |  |
| **Age** |  |  |  |  |
| ≤55 years | 6 (10.0) | 3 (15.8) | 3 (7.3) | 0.370 |
| >55 years | 54 (90.0) | 16 (84.2) | 38 (92.7) |  |
| **Gender** |  |  |  |  |
| Female | 21 (35.0) | 9 (47.4) | 1 (29.3) | 0.245 |
| Male | 39 (65.0) | 10 (52.6) | 29 (70.7) |  |
| **HPD** |  |  |  |  |
| Clear cell | 30 (50.0) | 8 (42.1) | 22 (53.7) | 0.354 |
| Papillary | 15 (25.0) | 4 (21.1) | 11 (26.8) |  |
| Chromophobic | 15 (25.0) | 7 (36.8) | 8 (19.5) |  |
| **Tumor location** |  |  |  |  |
| Right side | 22 (36.7) | 7 (36.8) | 15 (36.6) | 0.985 |
| Left side | 38 (63.3) | 12 (63.2) | 26 (63.4) |  |
| **Grade** |  |  |  |  |
| Grade 1 | 9 (15.0) | 1 (5.3) | 8 (19.5) | **0.023** |
| Grade 2 | 28 (46.7) | 6 (31.6) | 22 (53.7) |  |
| Grade 3 | 23 (38.3) | 12 (63.2) | 11 (26.8) |  |
| **Tumor size** |  |  |  |  |
| T1 | 21 (35.0) | 6 (31.6) | 15 (36.6) | 0.589 |
| T2 | 25 (41.7) | 7 (16.8) | 18 (43.9) |  |
| T3 | 14 (23.3) | 6 (31.6) | 8 (19.5) |  |
| **LN** |  |  |  |  |
| Negative | 39 (65.0) | 11 (57.9) | 28 (68.3) | 0.562 |
| Positive | 21 (35.0) | 8 (42.1) | 13 (31.7) |  |
| **Recurrence** |  |  |  |  |
| Negative | 44 (73.3) | 7 (36.8) | 37 (90.2) | **<0.001** |
| Positive | 16 (26.7) | 12 (63.2) | 4 (9.8) |  |
| **DFS** |  |  |  |  |
| ≤ 1year | 29 (48.3) | 19 (100) | 10 (24.4) | **<0.001** |
| > 1year | 31 (51.7) | 0 (0.0) | 31 (75.6) |  |
| **GAS5 fold** |  |  |  |  |
| ≤ 1-fold | 55 (91.7) | 19 (100) | 36 (87.8) | 0.283 |
| > 1-fold | 3 (5.0) | 0 (0.0) | 3 (7.3) |  |
| >10-folds | 2 (3.3) | 0 (0.0) | 2 (4.9) |  |

| **miR-34a fold** |  |  |  |  |
| --- | --- | --- | --- | --- |
| ≤ 1-fold | 15 (25.0) | 6 (31.6) | 9 (22.0) | 0.192 |
| > 1-fold | 19 (31.7) | 8 (42.1) | 11 (26.8) |  |
| >10-folds | 26 (43.3) | 5 (26.3) | 21 (51.2) |  |

*HPD* histopathological diagnosis, *DFS* disease-free survival, *OS* overall survival. Low OS: ≤12 months, High OS: >12 months. Fisher's Exact and Pearson Chi-square tests were used. Bold values indicate statistical significance at $p < 0.05$

**Table 2.** Clinicopathological characteristics of glioblastoma patients.

| **Variables** | **Total** | **Low OS** | **High OS** | ***p* value** |
| --- | --- | --- | --- | --- |
| **Total number** | 50 (100) | 10 (20.0) | 40 (80.0) |  |
| **Age** |  |  |  |  |
| ≤55 years | 21 (42.0) | 7 (70.0) | 14 (35.0) | 0.073 |
| >55 years | 29 (58.0) | 3 (3.0) | 26 (65.0) |  |
| **Gender** |  |  |  |  |
| Female | 13 (26.0) | 1 (10.0) | 12 (30.0) | 0.258 |
| Male | 37 (74.0) | 9 (90.0) | 28 (70.0) |  |
| **Tumor location** |  |  |  |  |
| Frontal | 24 (48.0) | 8 (80.0) | 16 (40.0) | 0.071 |
| Temporo-paroetal | 18 (36.0) | 1 (10.0) | 17 (42.5) |  |
| Fronto-temporal | 8 (16.0) | 1 (10.0) | 7 (17.5) |  |
| **Recurrence** |  |  |  |  |
| Negative | 44 (88.0) | 8 (80.0) | 36 (90.0) | 0.586 |
| Positive | 6 (12.0) | 2 (20.0) | 4 (10.0) |  |
| **DFS** |  |  |  |  |
| ≤ 1year | 25 (50.0) | 10 (100) | 15 (37.5) | **0.001** |
| > 1year | 25 (50.0) | 0 (0.0) | 25 (62.5) |  |
| **GAS5 fold** |  |  |  |  |
| ≤ 1-fold | 37 (74.0) | 7 (70.0) | 30 (75.0) | 0.707 |
| > 1-fold | 13 (26.0) | 3 (30.0) | 10 (25.0) |  |
| >10-folds |  |  |  |  |
| **miR-34a fold** |  |  |  |  |
| ≤ 1-fold | 38 (76.0) | 8 980.0) | 30 (75.0) | 0.741 |
| > 1-fold | 12 (24.0) | 2 (20.0) | 10 (25.0) |  |
| >10-folds | 0 (0.0) |  |  |  |

Data are presented as number (frequency). *HPD* histopathological diagnosis, *DFS* disease-free survival, *OS* overall survival. Low OS: ≤12 months, High OS: >12 months. Fisher's Exact and Pearson Chi-square tests were used. Bold values indicate statistical significance at $p < 0.05$

**Table 3.**
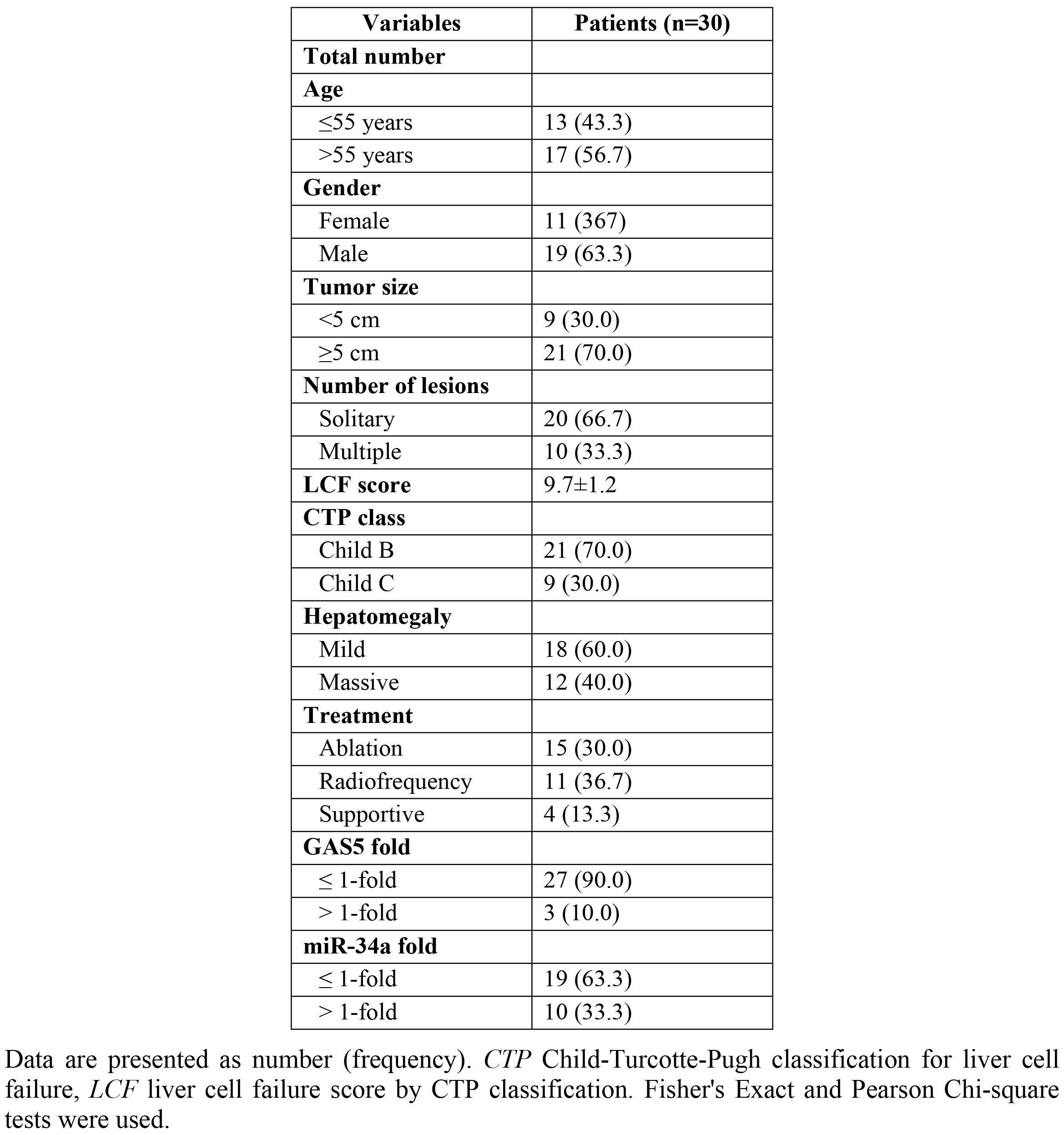
Clinicopathological characteristics of hepatocellular carcinoma patients.

| Variables | Patients (n=30) |
| --- | --- |
| **Total number** |  |
| **Age** |  |
| ≤55 years | 13 (43.3) |
| >55 years | 17 (56.7) |
| **Gender** |  |
| Female | 11 (36.7) |
| Male | 19 (63.3) |
| **Tumor size** |  |
| <5 cm | 9 (30.0) |
| ≥5 cm | 21 (70.0) |
| **Number of lesions** |  |
| Solitary | 20 (66.7) |
| Multiple | 10 (33.3) |
| **LCF score** | 9.7±1.2 |
| **CTP class** |  |
| Child B | 21 (70.0) |
| Child C | 9 (30.0) |
| **Hepatomegaly** |  |
| Mild | 18 (60.0) |
| Massive | 12 (40.0) |
| **Treatment** |  |
| Ablation | 15 (30.0) |
| Radiofrequency | 11 (36.7) |
| Supportive | 4 (13.3) |
| **GAS5 fold** |  |
| ≤ 1-fold | 27 (90.0) |
| > 1-fold | 3 (10.0) |
| **miR-34a fold** |  |
| ≤ 1-fold | 19 (63.3) |
| > 1-fold | 10 (33.3) |

Data are presented as number (frequency). *CTP* Child-Turcotte-Pugh classification for liver cell failure, *LCF* liver cell failure score by CTP classification. Fisher's Exact and Pearson Chi-square tests were used.

### Expression profiling

GAS5 levels were under-expressed in RCC [0.08 (0.006-0.38), *p* <0.001], GB [0.10 (0.003-0.89), *p* < 0.001] and HCC [0.12 (0.015-0.74), *p* < 0.001]. On the other hand, miR-34a displayed an increased expression in RCC [4.05 (1.003-22.69), *p* < 0.001] and a decreased expression in GB [0.35 (0.04-0.95), *p* < 0.001] as depicted in Fig. 3. A weak negative correlation was observed between levels of GAS5 and miR-34a in GB [r = −0.39, *p* = 0.006] (Fig. 4).

**Fig. 3.**
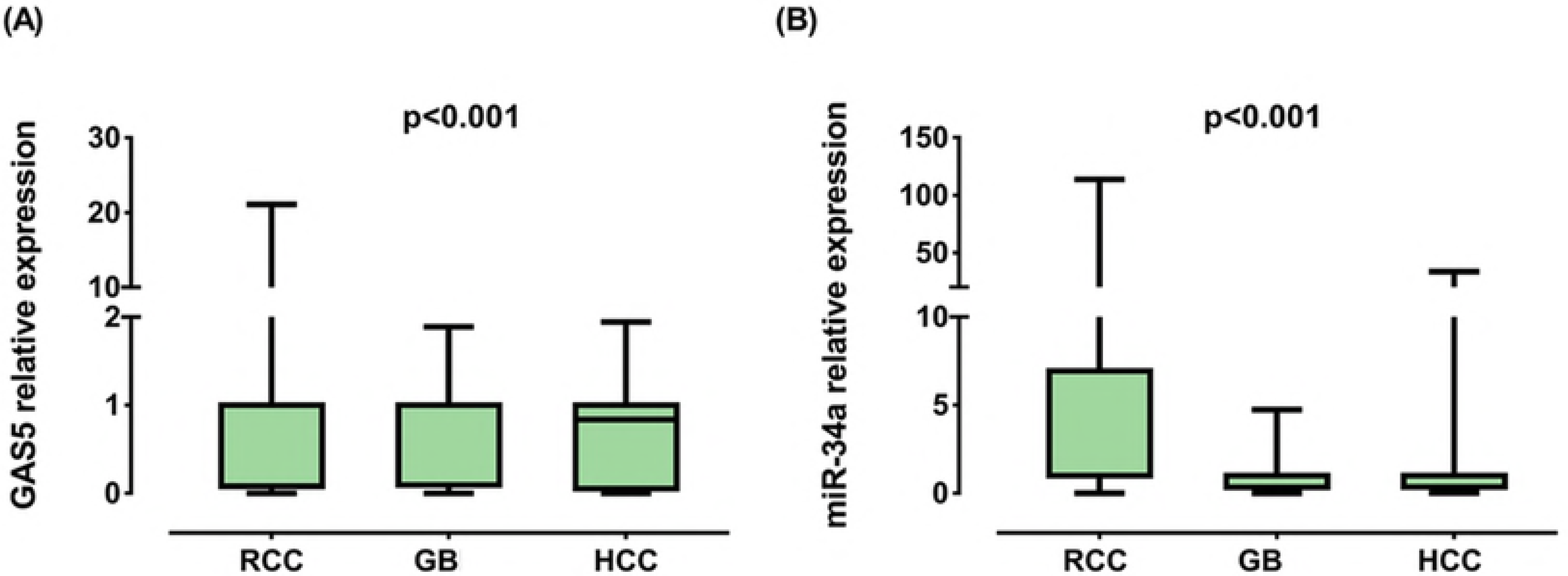
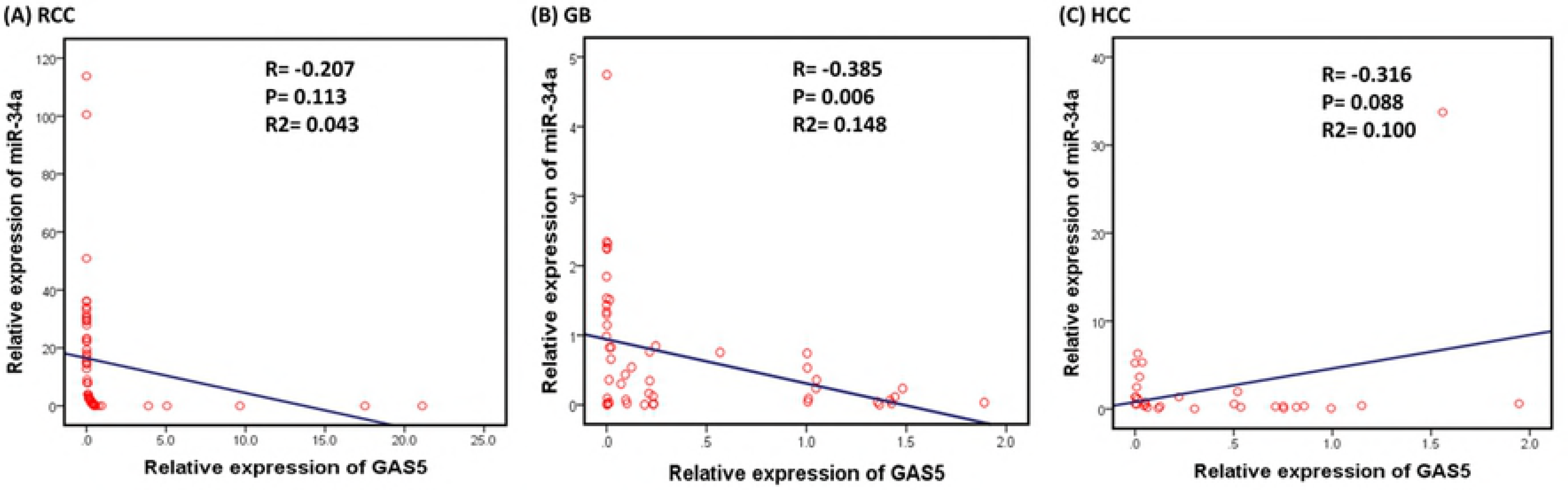
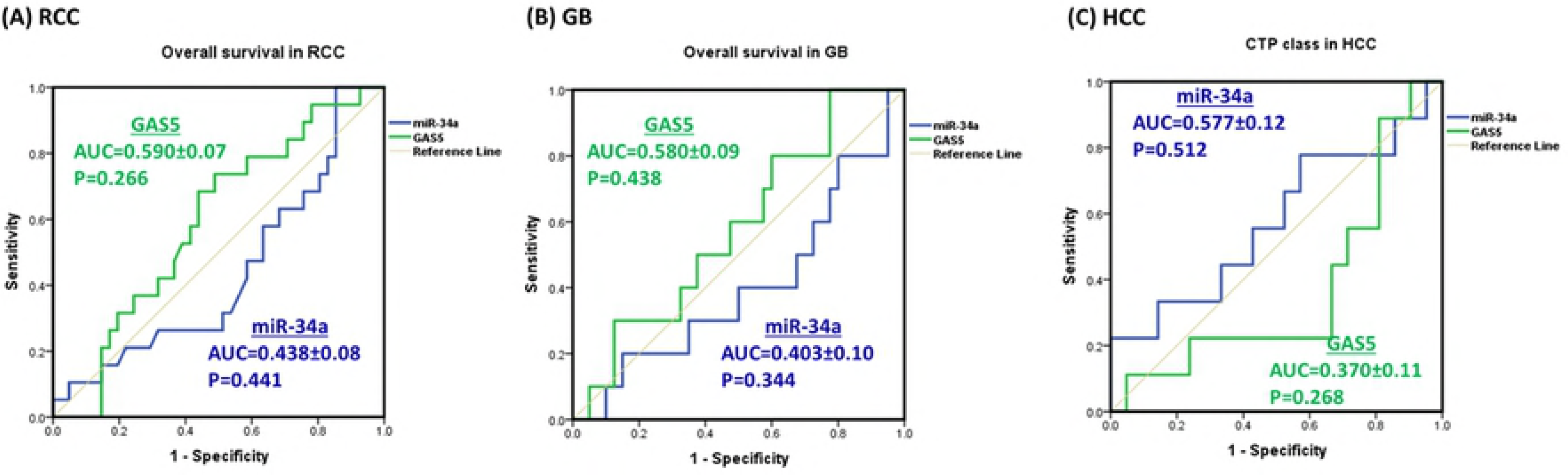
Relative expression of GAS5 and miR-34a-5p in cancer. *RCC* renal cell carcinoma, GB glioblastoma, *HCC* hepatocellular carcinoma. Data are represented as medians. The box defines upper and lower quartiles (25% and 75%, respectively) and the error bars indicate upper and lower adjacent limits. Expression levels in cancer and control samples were normalized to GAPDH in RCC and HCC, TBP in GB and RNU6B for microRNA and calculated using the delta-delta CT method [= 2 (−ΔΔCT)] in comparison to controls. Fold change of controls were set at 1.0. Wilcoxon matched-pair signed-rank and Mann-Whitney U tests were used for tissues of renal/brain cancer and serum of liver cancer patients, respectively. Two-sided *p* < 0.05 was considered statistically significant

**Fig. 4.**
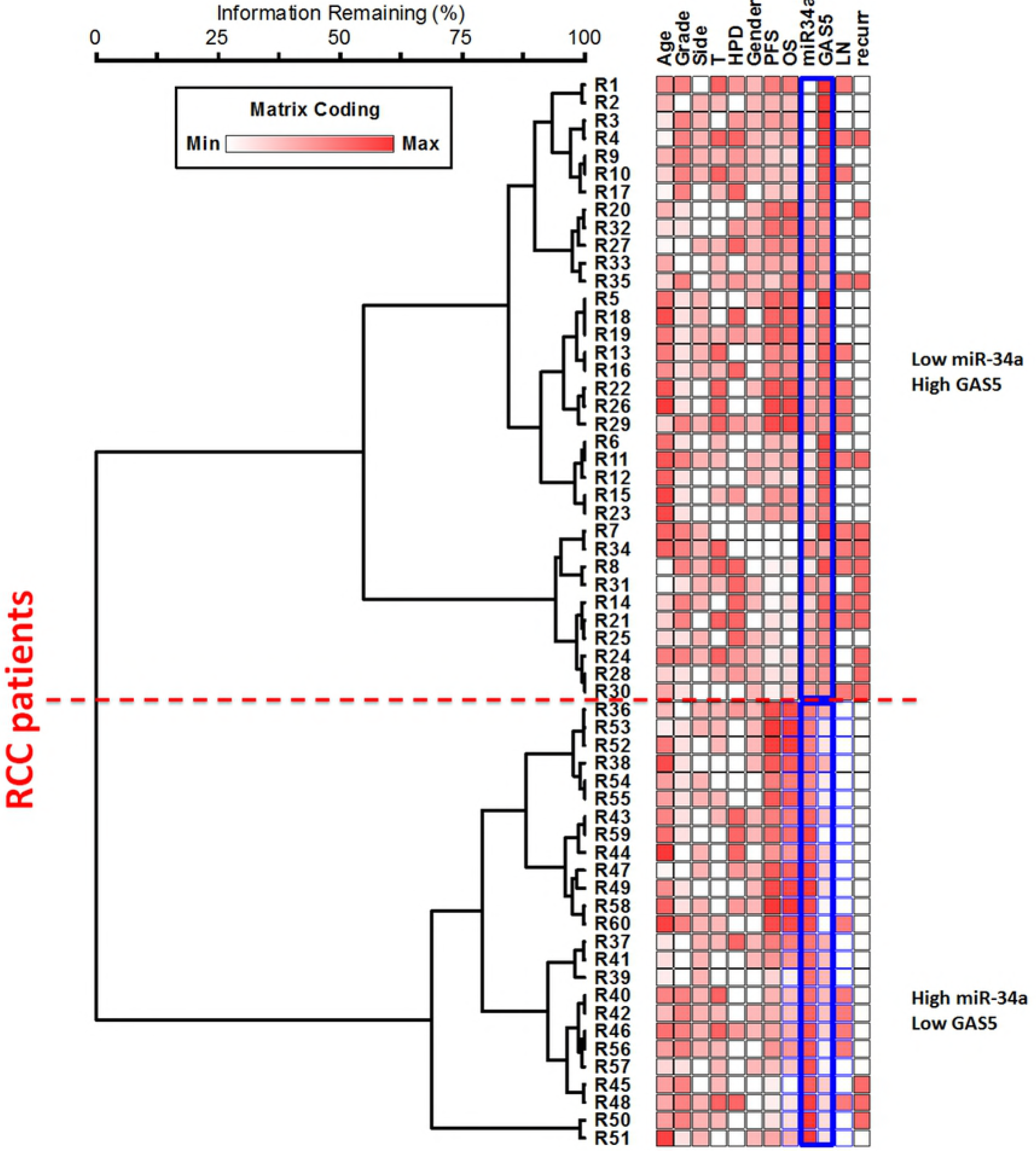
Correlation between GAS5 and miR-34a-5p expression levels. *RCC* renal cell carcinoma, *GB* glielation test was used. Statistical significance was considered at *p* < 0.0oblastoma, *HCC* hepatocellular carcinoma. Spearman’s rank corr5.

### Association of GAS5 and miR-34a with clinicopathological features

Univariate analyses are shown in Table 4A, B, and C. The lower GAS5 level was correlated with poor DFS (r = 0.31, *p* = 0.018) and OS (r = 0.28, *p* = 0.029) in RCC but not in GB, and was correlated by a marginal significance (r= −0.35, *p* = 0.056) with a higher number of lesions in HCC. Levels of miR-34a did not correlate with any clinico-pathological features.

**Table 4.**
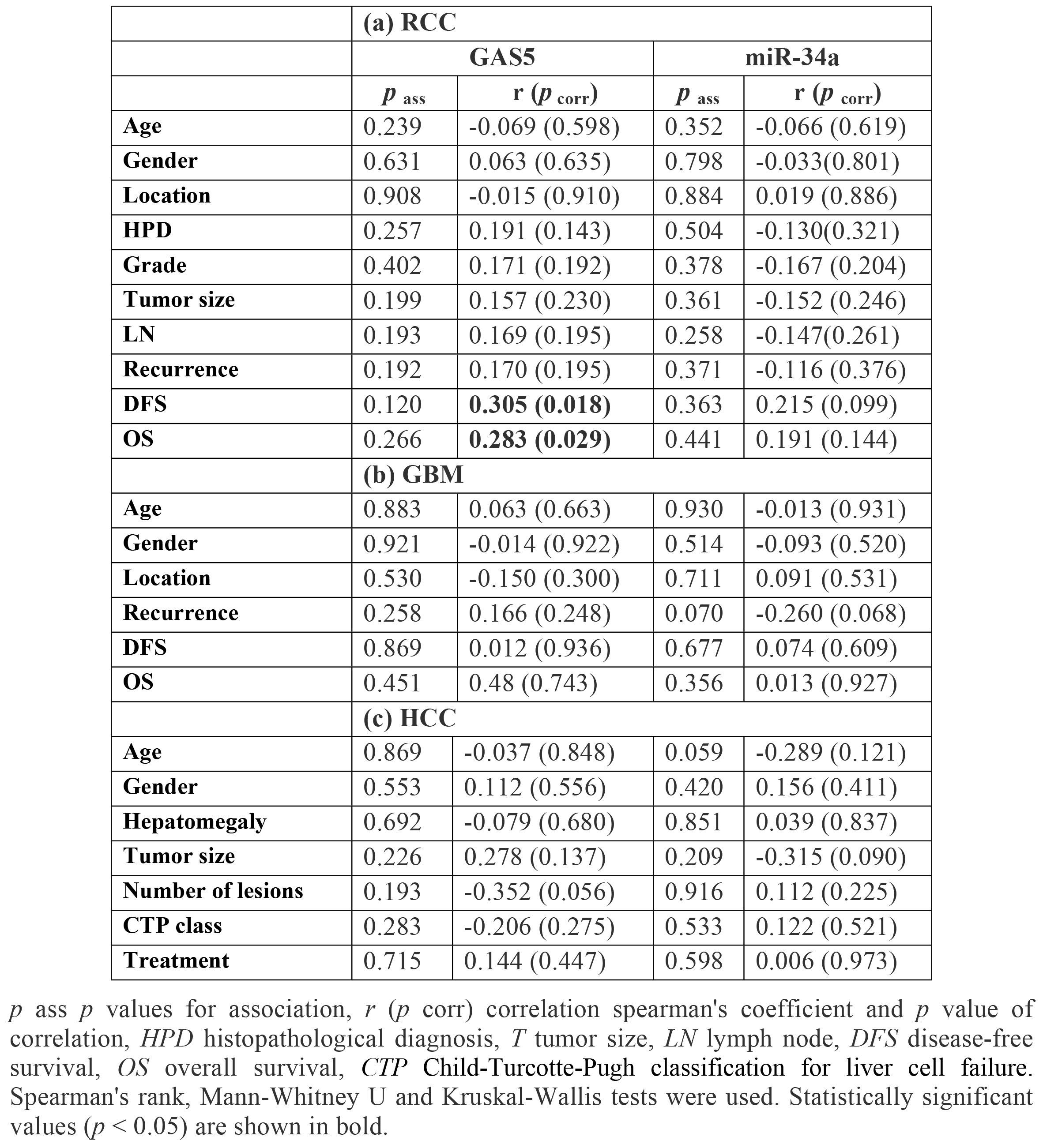
Univariate analysis for association between gene profile and clinico-pathological features in the study cohorts.

### Role of GAS5 and miR-34a as markers of cancer prognosis

ROC curves showed no prognostic value for GAS5 nor miR-34a to predict survival of cancer patients in RCC and GB, or to predict CTP class for liver cell failure in HCC patients (*p* > 0.05) (Fig. 5).

**Fig. 5.**
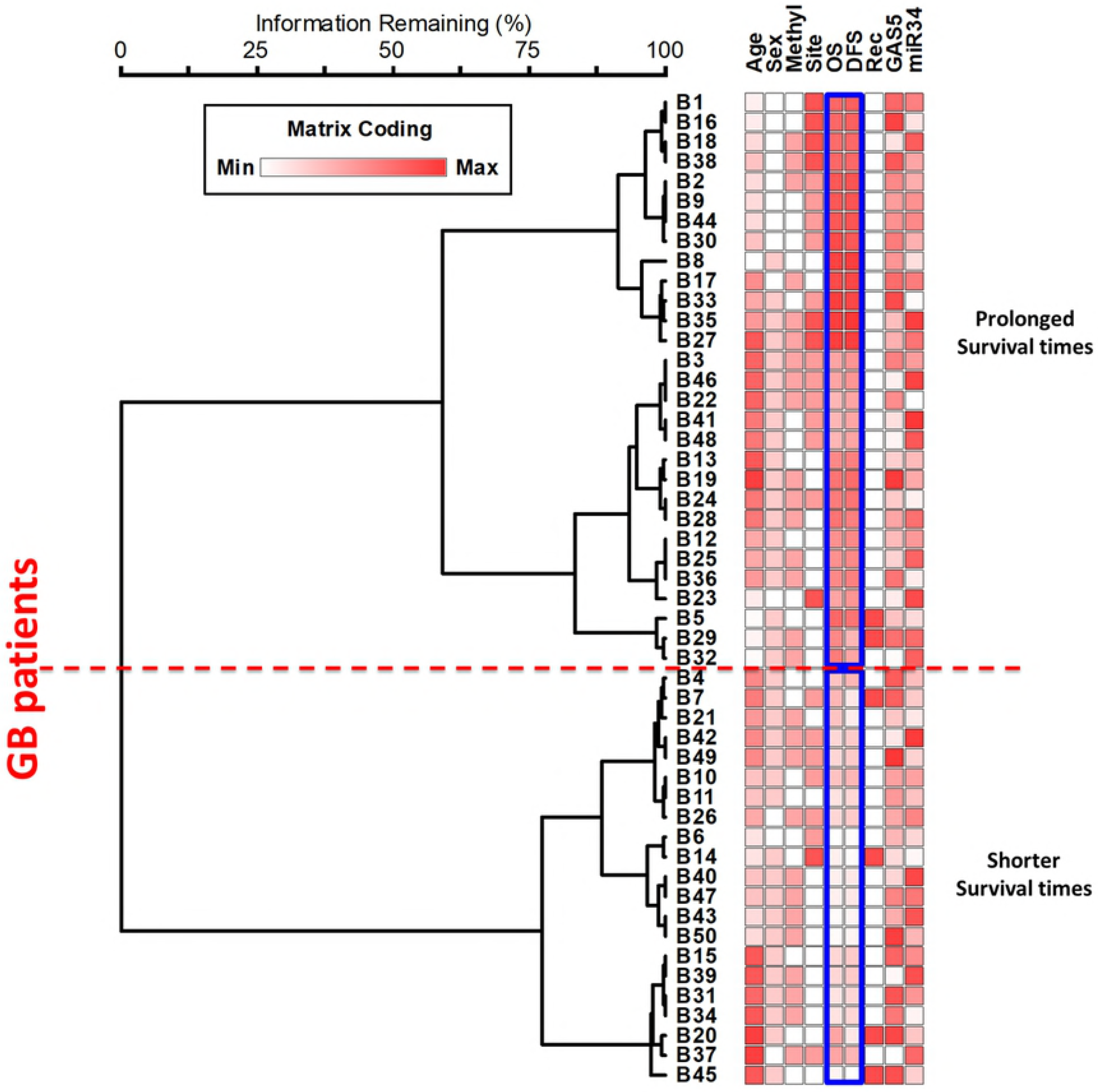
Receiver Operating Characteristics (ROC) analysis for the prognostic value of markers. *RCC* renal cell carcinoma, *GB* glioblastoma, *HCC* hepatocellular carcinoma, *AUC* area under curve, *CTP* Child-Turcotte-Pugh classification for liver cell failure

### Survival analysis in RCC and GB

In RCC, multivariable analysis using logistic regression test (Enter method) showed age and pathological grade to be independent predictors for recurrence: (OR = 1.251, 95% CI = 1.075-1.455, *p* = 0.004) for the age and (OR = 19.9, 95% CI = 1.034-383, *p* = 0.047) for the grade. Kaplan-Meier curve analysis and log-rank test revealed that RCC patients with female gender, postnephrectomy recurrence, advanced pathological grade, and down-regulated miR-34a levels had significantly poor overall survival than their corresponding (*p* < 0.05) (Table 5). In addition, multivariable analysis by Cox regression model demonstrated gender and recurrence to be independent predictors of overall survival (hazard ratio (HR) = 2.49; 95 % confidence interval (95 % CI) 1.14–5.41, *p* = 0.021) and (HR = 4.16; 95% CI of 1.88–9.16, *p* < 0.001), respectively.

**Table 5.** Multivariable analysis in renal cell carcinoma patients.

| Variables | Survival time | Overall comparisons |  |  | Cox regression |  |
| --- | --- | --- | --- | --- | --- | --- |
|  | OS (mo) | Log Rank | Breslow | Tarone-Ware | HR (95% CI) | *p* |
| **Age** |  |  |  |  |  |  |
| ≤55 years | 13.1 ± 2.22 |  |  |  |  |  |
| >55 years | 16.1 ± 0.8 | 0.186 | 0.244 | 0.221 | 0.97 (0.92-1.01) | 0.217 |
| **Gender** |  |  |  |  |  |  |
| Female | 13.9 ± 1.28 |  |  |  |  |  |
| Male | 16.7 ± 0.94 | **0.027** | 0.077 | 0.095 | **2.49 (1.14-5.41)** | **0.021** |
| **HPD** |  |  |  |  |  |  |
| Type 1 | 16.1 ± 1.12 |  |  |  |  |  |
| Type 2 | 17.5 ± 1.61 |  |  |  | 0.70 (0.34-1.46) | 0.351 |
| Type 3 | 13.3 ± 1.25 | 0.103 | 0.198 | 0.159 | 0.69 (0.30-1.60) | 0.397 |
| **Location** |  |  |  |  |  |  |
| Right side | 16.3 ± 1.23 |  |  |  |  |  |
| Left side | 15.4 ± 1.00 | 0.703 | 0.489 | 0.539 | 1.08 (0.57-2.06) | 0.807 |
| **Grade** |  |  |  |  |  |  |
| Grade 1 | 16.3 ± 1.53 |  |  |  |  |  |
| Grade 2 | 18.0 ± 1.17 |  |  |  |  |  |
| Grade 3 | 12.8 ± 1.05 | **0.005** | **0.003** | **0.003** | 1.60 (0.66-3.85) | 0.293 |
| **Tumor size** |  |  |  |  |  |  |
| T1 | 16.4 ± 1.40 |  |  |  |  |  |
| T2 | 15.9 ± 1.13 |  |  |  | 0.91 (0.30-2.75) | 0.871 |
| T3 | 14.5 ± 1.65 | 0.709 | 0.577 | 0.635 | 0.89 (0.34-2.29) | 0.813 |
| **LN** |  |  |  |  |  |  |
| Negative | 16.4 ± 0.97 |  |  |  |  |  |
| Positive | 14.6 ± 1.27 | 0.306 | 0.235 | 0.255 | 1.05 (0.42-2.59) | 0.911 |
| **Recurrence** |  |  |  |  |  |  |
| Negative | 17.5 ± 0.83 |  |  |  |  |  |
| Positive | 10.8 ± 1.07 | **<0.001** | **<0.001** | **<0.001** | **4.16 (1.88-9.16)** | **<0.001** |
| **GAS5 fold** |  |  |  |  |  |  |
| Under-expressed | 15.7 ± 0.84 |  |  |  |  |  |
| Over-expressed | 16.2 ± 1.24 | 0.735 | 0.753 | 0.985 | 0.29 (1.04-0.96) | 0.291 |
| **miR-34a fold** |  |  |  |  |  |  |
| Under-expressed | 13.2 ± 0.99 |  |  |  |  |  |
| Over-expressed | 16.6 ± 0.95 | **0.010** | **0.049** | **0.024** | 1.001 (0.98-1.01) | 0.926 |

Survival times is shown as mean and standard error, *OS* overall survival, *HR (95% CI)* Hazard ratio (95% confidence interval), *HPD* histopathological diagnosis, *T* tumor size, *LN* lymph node,. Statistically significant values ( $p < 0.05$ ) are shown in bold.

In GB, Kaplan-Meier curves showed a significant association of shorter survival times with male gender (Breslow test: *p* = 0.002 and Tarane-Ware test: *p* = 0.030). In addition, marginal significance was observed for poor overall survival in patients with frontal lesions (Log rank test: *p* = 0.050) (Table 6). Multivariable analysis by Cox regression model illustrated tumor recurrence to be an independent predictor of low overall survival (HZ = 11.1; 95 % CI 2.88-42.5, *p* < 0.001).

**Table 6.**
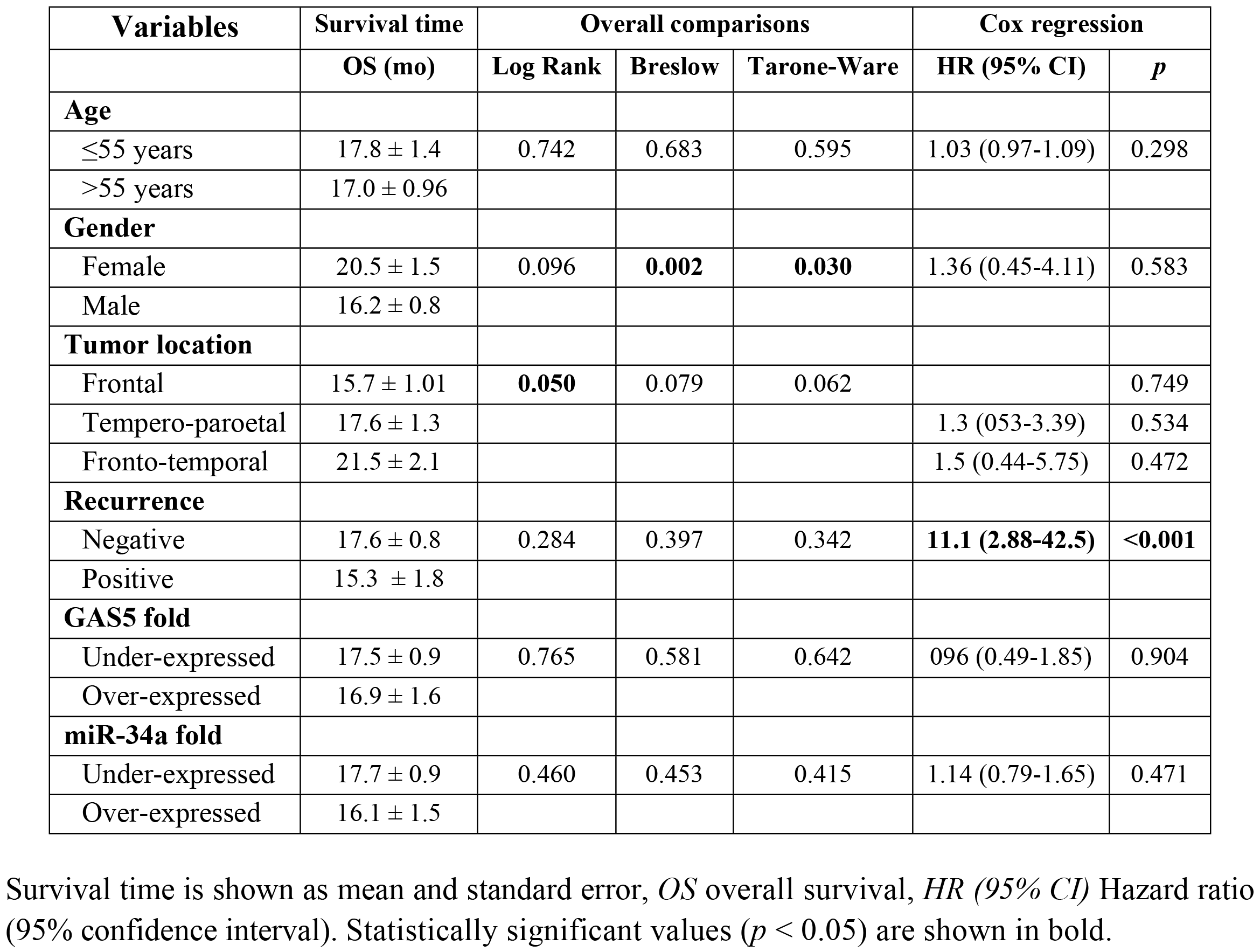
Multivariable analysis in glioblastoma patients.

### Hierarchical clustering analysis

Dendrograms for Two-way agglomerative hierarchical cluster analysis, were employed (Fig. 6). The following cluster setup parameters were adjusted: Distance method: Sorensen (Bray-Curtis), Group Linkage Method: Flexible Beta at 0.75, Clustering of factor relative by factor maximum. A distance matrix is shown. RCC plot analyzed 60 strands and 12 factors, GB plot analyzed 50 strands and 9 factors, whereas HCC dendrogram showed results of 30 strands and 11 factors. Clustering analysis revealed separation of RCC patients by GAS5 and miR-34a levels, GB patients by survival times, and HCC patients by their age.

**Fig. 6.**
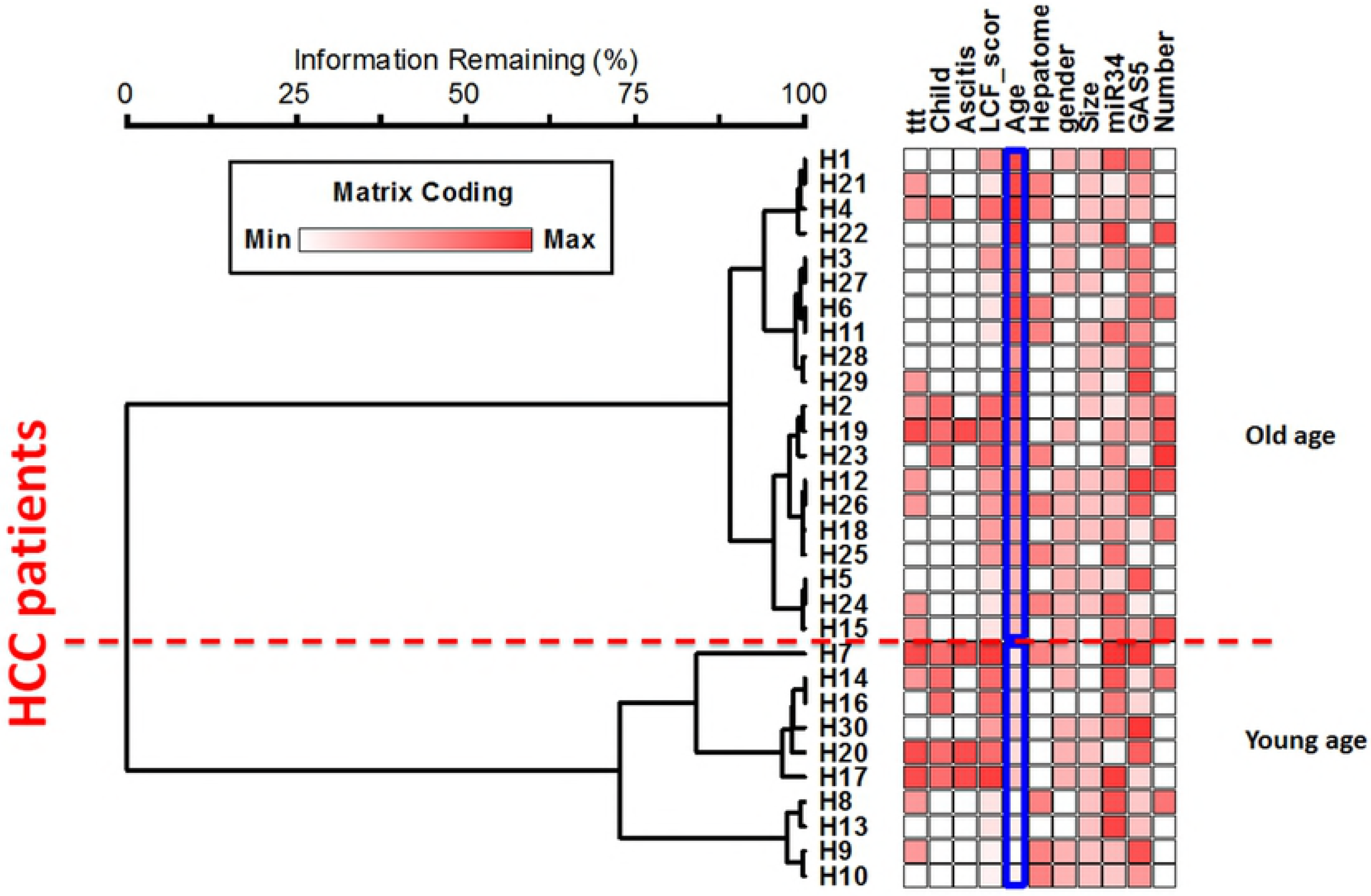
Multivariate analysis of patients according to combined transcriptomic signature of genes and clinicopathological features. (A) RCC, (B) GB, (C) HCC. RCC patients were divided based on GAS5 and miR-34a levels, GB patients were divided by survival times, and HCC patients by age

## Discussion

In this study, we measured the expression of two ncRNAs; the lncRNA GAS5 and the miRNA miR-34a in three of the most prevalent and high-incidence tumors in Egypt; hepatic, renal and brain cancer. We chose more than one tumor type to assess if the same ncRNA could work differently according to the tissue type. We also investigated the possible association between GAS5 and miR-34a in mediating carcinogenesis after detecting an interaction between the two through our preliminary in *silico* analysis.

Our results show that *GAS5* was under-expressed in the three types of cancer; RCC, HCC and GB. On the other hand, levels of miR-34a greatly varied according to the tumor type. RCC patients had a lower GAS5 level and a higher miR-34a level in tumor tissue compared to adjacent normal tissue. Moreover, GAS5 levels were correlated with poor survival. In accordance, Qiao et al. reported a reduced GAS5 level in RCC cell lines compared to normal cell lines. Furthermore, in *vitro* cloning and functional expression analysis revealed that *GAS5* overexpression caused cell cycle arrest, increased apoptosis, as well as inhibited tumor metastatic potential [24], which could explain the correlation between reduced patient survival and GAS5 level in our study. As for miR-34a, several studies reported over-expression of this RNA in RCC patients [40, 41, 54–56]. Liu et al. predicted that its oncogenic function could be through targeting the two tumor suppressors secreted frizzled related protein 1 (SFRP1) and calmodulin binding transcription activator 1 (CAMTA1), and further validated the first target [40]. Another study noticed that miR-34a enhances cell survival, both in *vitro* and in *vivo* during cisplatin nephrotoxicity through p53 [57]. Collectively, the previous studies suggest that miR-34a can act as an oncogene in renal tissue. On the contrary, Yadav et al. detected a significant under-expression of miR-34a in both sera and renal tissues of RCC patients [58]. Similar results were demonstrated by the studies of Zhang et al. and Weng et al. [59, 60], where the former study demonstrated that decreased expression of miR-34a in RCC patients inversely up-regulated the gene for the transcription factor YY1. The latter study suggested that underexpression of miR-34a in cancer tissues of RCC patients affected the regulation of the *NOTCH1* gene, as well as caused dysregulation of other multiple miR-34a targets in 786-O and Caki-1 RCC cell lines. Yu et al. found that miR-34a suppressed tumor growth and metastasis in *vivo* and in *vitro* through targeting CD44 [61]. Another study found that miR-34a inhibits cellular invasion in renal cancer cell lines A498 and 769P through targeting the 3’ untranslated region (UTR) of the *c-myc* oncogene [62]. Also, the lncRNA NEAT1 (nuclear paraspeckle assembly transcript 1) was found to sponge miR-34a releasing its inhibition on the *c-met* oncogene, resulting in increased cellular proliferation and invasion in RCC cell lines 786-O and ACHN [63]. Furthermore, Bai et al. found that miR-34a causes senescence of rat renal cells through targeting two anti-oxidative mitochondrial genes [64]. Cellular senescence, which is an irreversible state of growth arrest, protects the cells from accumulating mutations that could lead to malignant transformation [65]. Further supporting the tumor suppressor potential of miR-34a, Zhou et al. provided evidence that miR-34a, secreted by fibroblasts, enhances apoptosis of renal tubular cells through regulating the anti-apoptotic gene *BCL-2* [66].

The inconsistent results between our study which shows up-regulation of *miR-34a* in RCC and other studies showing its under-regulation in the same type of cancer could have many possible explanations. First, we measured the expression of miR-34a in cancer tissues obtained directly from RCC patients as opposed to other studies conducted on cancer cell lines. While those cell lines are highly essential for functional molecular analysis, they may be different from primary tumors, possibly through building up new mutations in their attempt to adjust to their artificial environment [67, 68]. Such mutations may easily alter cellular responses and regulatory mechanisms, possibly affecting the expression of *miR-34a*. Second, miR-34a could be a non-specific molecule that can both activate or inhibit tumerogenesis depending on the surrounding environment. These include internal stimuli (other regulatory molecules or polymorphisms, oxidative molecules, other associated disease states, tumor stage/grade or else) and external stimuli (cellular response to environmental exposures, including chemotherapy, other drugs, chemicals, foods, etc). Third, miR-34a has multiple targets (discussed in our previous work) [41], as well as being itself a target for many lncRNAs. Each study focuses on one or few targets, and infers a tumor suppressor or an oncogenic function based on its effect on the studied target/pathway. However, miR-34a is one small molecule in a larger network of molecules that either promotes or inhibits tumorogenesis based on the net result of *all* its regulated targets, which could easily differ according to countless variables. Given by the negative correlation of this microRNA with GAS5 in our RCC patients, as well as the predicted interaction between the two, we believe that a new pathway; the *GAS5*/miR-34a pathway might be involved in the previously indicated molecular network leading to RCC.

The interaction between lncRNAs and miRNAs can be multifaceted. Yoon et al. explained four mechanisms of interaction between the two types [69]. First, lncRNAs can act as miRNA sponges as previously mentioned with examples in the introduction. Second, the opposite can occur, where miRNAs can inhibit lncRNAs by binding to them and causing their degradation. This applies for miR145-5p, miR-181a-5p and miR-99b-3p which inhibit lncRNA ROR [70], and miR-9 which inhibits lncRNA MALAT1 [71]. Third, miRNAs and lncRNAs can both compete for the same binding site on mRNAs, an example of which is miR-485-5p and lncRNA BACE1AS competing for BACE1 mRNA [72]. Fourth and finally, another form of relationship exists between the two types; where the lncRNA (>200 nucleotide) is capable of generating smaller (<22 nucleotide) miRNAs such as the lncRNA H19 generating miR-675 [73]. GAS5 provides a perfect example for such interactions. For instance, GAS5 acts as a molecular sponge for many different microRNAs, including miR-21, miR-222, miR-196a, miR-205, miR-221 and miR-103. [16, 26, 74–77], all of which are related to cancer. Then again, both miR-21 and miR-222 can negatively regulate *GAS5* [27, 78]. Also, three of the snoRNAs produced by GAS5 (U44, U74 and U78) can give rise to miRNAs [79], making GAS5 one of the lncRNAs generating miRNAs.

While functional validation has been yet required to prove the direct interaction between GAS5 and miR-34a, it is highly plausible. This is due to the already verified interaction between miR-34a and GAS1 (Growth arrest specific 1), another member of the *GAS* genes [42]. *GAS1* is a protein coding gene that, similar to *GAS5*, exerts its tumor suppressor actions through arresting the cell cycle and stimulating apoptosis [80]. Ma et al. measured the expression of miR-34a and the GAS1 protein in papillary thyroid carcinoma; GAS1 was under-expressed, while miR-34a was over-expressed. Further analysis revealed that miR-34a binds to the 3’UTR of GAS1 causing its silencing, which in turn activates the *RET* oncogene [42]. Through the BLAST tool, we detected sequence homology between the *GAS1* and the *GAS5* genes, further raising the possibility of interaction of miRNA-34a with GAS5. Given that *GAS5* is down-regulated and *miR-34a* is up-regulated in our study, tumorogenesis may be in such a case due to sponging of miR-34a by GAS5, where underexpression of *GAS5* in the rapidly dividing cancer cells causes release of the inhibition of miR-34a on its tumor suppressor targets, allowing for further tumor progression. Oppositely, the inverse correlation between the two could also suggest that miR-34a inhibits GAS5, making its underexpression a cause of cancer rather than a result, as in the case of the miR-34a-GAS1 interaction. miR-34a expression levels in our study were not significant in the case of patients with HCC. *GAS5* levels, however, were under-expressed in HCC patients. In addition, lower GAS5 levels were associated with more numerous tumor foci. The same conclusions were reached by studies conducted by Tu et al., Chang et al. and Hu et al. who found that *GAS5* was under-expressed in HCC patients and predicted poor survival in those patients [15, 26, 81]. The latter study by Hu et al. found that under-expression of *GAS5* in HCC cell lines releases its sponging effect on oncomiR-21, which normally targets the two tumor suppressor genes *PDCD4* and *PTEN* [26]. On the contrary, according to Tao et al., *GAS5* was a proto-oncogene in HCC where they found an indel polymorphism in the promotor of *GAS5* that increased the risk of HCC in Chinese. In *vitro* analysis showed that the deletion allele of the polymorphism altered methylation of the *GAS5* promoter and was associated with higher levels of GAS5 in HCC cell lines Sk-hep-1, Bel-7404 and Huh7. Further analysis provided evidence that the resulting over-expression of *GAS5* had an anti-apoptotic effect in those cell lines [82]. This divergence observed by Tao et al. could be due to racial differences. In other words, the allele causing *GAS5* over-expression might be more common in their studied population. To further confirm this probability, we searched the 1000 genome project phase 3 databases [83] for this indel polymorphism (positon 1:173868254-173868258 (AGGCA/-)). We found that the frequency of the deletion allele was very high in Asians and Chinese in particular (28-40%) as compared to other races (3-12%), showing that in other populations the effect of the polymorphism on HCC and *GAS5* expression could be negligible.

In patients with GB, both GAS5 and miR-34a levels were significantly down-regulated. Zhang et al. correlated the expression of five lncRNAs including GAS5 with GB, and found that higher levels of GAS5 were associated with prolonged survival [84]. Zhao et al. found that *GAS5* was under-expressed in glioma cell lines U87 and U251 and that its tumor suppressor role was through targeting miR-222 [16]. The same author, in a more recent study, added another miRNA to the targets of GAS5; miR-196a-5p, where *GAS5* under-expression in human glioma stem cells enhanced tumor progression through inhibiting this miRNA [85]. Regarding the more controversial *miR-34a*, studies on brain cancer, including GB and glioma agree on its role as a tumor suppressor in this particular tissue; reviewed in [86], which fits in with its expression status in the present study.

## Conclusions

In this study, we show that *GAS5* is under-expressed in three types of tumors, in addition to being associated with tumor prognosis in some types. Consequently, we believe that GAS5 could potentially be used as a prognostic marker for cancer. Furthermore, in favor of this notion is the predicted subcellular localization of GAS5 provided by our in *silico* analysis, which shows that GAS5 is most significantly localized in the extracellular exosomes, supporting its existence in the circulation and highlighting its putative role as a non-invasive biomarker which mirrors tissue pathology. Furthermore, we suppose that miR-34a might be a potential target of GAS5, or *vice versa*. However, we confirm that further studies are required to validate this interaction.

## Acknowledgements

The authors thank the Center of Excellence in Molecular and Cellular Medicine and the Oncology Diagnostic Unit, Suez Canal University, Ismailia, Egypt for providing the facilities for performing the research work.

## Compliance with ethical standards

### Conflicts of interest

The authors declare that they have no competing interests.

### Funding

None.

## Supporting Information

**S1 Table: GAS5 alternative splicing transcripts.**

**S2 Table: GAS5 common variants.**

**S3 Table: LncRNA GAS5-microRNA interaction.**

**S4 Table: Complementarity between GAS5 transcripts and miR-34a-5p.**

**S5 Table: hsa-miR-34a pathways.**

